# Perceiving direction of deformation-based motion

**DOI:** 10.1101/2024.12.25.630338

**Authors:** Takahiro Kawabe

## Abstract

Transparent liquid flow deforms its background through light refraction, creating complex spatiotemporal patterns that challenge the visual system in discerning flow direction. This study investigates how the visual system perceptually resolves the direction of transparent liquid flow. Using two-dimensional 1/f noise as background, we sequentially deformed the noise with pre-defined displacement maps displaced across frames, systematically manipulating spatial frequency and displacement magnitude. Participants reported the perceived direction of deformation-based motion. The results emphasize the critical role of the relationship between the minimum wavelength and displacement magnitude. Performance declined when the displacement magnitude exceeded a certain proportion relative to the wavelength. Conversely, performance also dropped significantly under conditions of small displacements combined with low cut-off frequencies. Further analysis revealed the local deformation possibly hinders the detection of deformation-based motion. These findings enhance understanding of how global and local motion cues interact, providing insight into the visual system’s processing of deformation-induced motion and transparency.

## Introduction

The visual system is capable of perceiving layers of surfaces by utilizing various cues. For instance, spatial luminance relationships serve as critical indicators for perceptual transparency^1,2^ . Additionally, image blur^3,4^ , differences in spatial frequency within overlapping texture patterns^5–7^ , and variations in motion direction or velocity along the same line of sight^8^ are also known to play essential roles.

In addition to these cues, recent studies^9–11^ have proposed that image deformation serves as a cue to transparency. Previous research demonstrated that applying image deformations with specific spatiotemporal frequency characteristics to static images can induce the perception of a transparent layer overlaying the static image. Such deformation-based transparency is frequently observed in transparent liquid flows^12,13^ . As transparent liquids flow, the image information of objects or scenes behind them undergoes continuous spatiotemporal deformation due to refraction. The visual system likely interprets these refraction-induced deformations as arising not from the objects or scenes themselves, but from deformations mediated by a transparent material, resulting in the perception of a transparent layer.

Transparent liquids are ubiquitous in natural and artificial environments, yet the mechanisms underlying their perceived motion remain poorly understood, limiting our comprehension of how the visual system processes deformation-based motion. Investigating the mechanisms underlying the perception of liquid flow direction may provide valuable insights into deformation-based transparency perception.

In this study, we investigated the spatial characteristics of direction perception in deformation-based motion (Movie 1). In our experiment, a static two-dimensional 1/f noise image was deformed using a deformation map that spatially warped the image’s pixels (Figure 1). The deformation map was linearly displaced across frames, resulting in a consistent spatial displacement of image deformation. We define deformation-based motion as the linear spatial shift of image deformation, driven by the spatial displacement of deformation maps. In contrast, we define “local deformation” as image deformation that can be generated by a single, static deformation map. Participants were tasked with reporting the perceived motion direction. The spatial frequency bandwidth and displacement magnitude of the deformation map were systematically manipulated. We aimed to investigate the interaction between the cut-off frequency of the spatial frequency bandwidth and the displacement magnitude in the discrimination of deformation-based motion direction. In a previous study using band-pass noise, it is known that the threshold for determining the direction of displacement depends on the spatial frequency of the band-pass noise^14^ . We hypothesized that a similar phenomenon may also be observed between displacement magnitude and the spatial frequency of image deformation. Specifically, we predicted that discrimination performance would decline when the displacement magnitude exceeds a certain proportion relative to the wavelength. Moreover, we also checked the interaction between the deformation-based motion and local deformation. Our results suggest that the detection of unidirectional motion signals plays a crucial role in judging the direction of deformation-based motion. Furthermore, we discuss the potential involvement of such unidirectional flows in the mechanisms underlying transparency perception.

**Figure 1.**
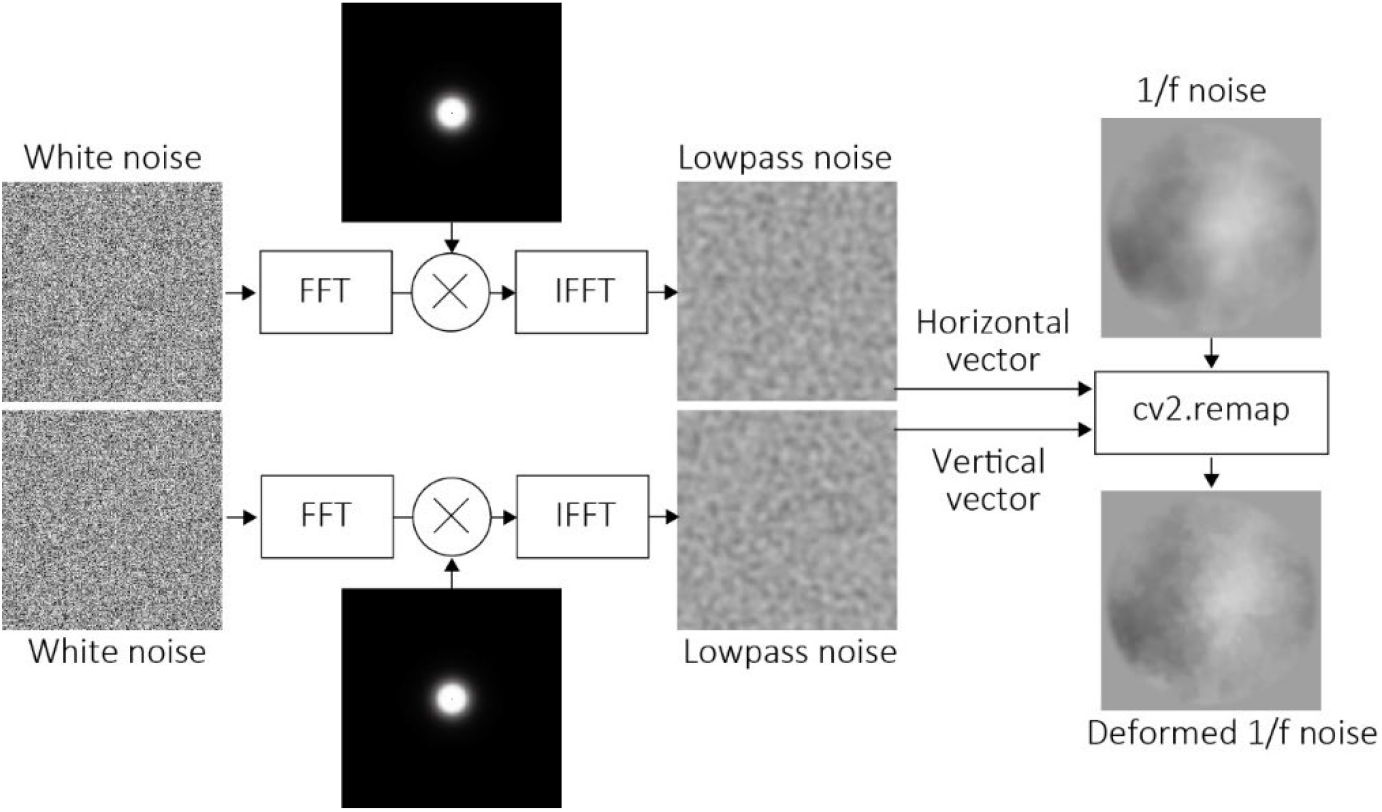
Schematic depiction of the stimulus generation process. Two sets of white noise were lowpass-filtered (in Experiment 1) or bandpass-filtered (in Experiments 2 and 3) and used as horizontal and vertical vectors to deform 1/f noise as the background. The deformed 1/f noise served as one of the video frames in a stimulus clip. The deformation displacement in the clip was generated by shifting the white noise.

## Method

### Participants

In Experiment 1, sixteen people (8 females) participated. Their mean age and standard deviation are 27.61 and 9.21, respectively. In Experiments 2, sixteen people (8 female) participated. Eight of the people had also participated in Experiment 1. Their mean age and standard deviation are 30.25 and 11.22, respectively. In Experiment 3, the people, who had participated in Experiment 2, also participated. Participants were recruited through a human resource agency in Japan and received monetary compensation based on criteria determined by the agency, which were not disclosed to the researchers. All participants were naive to the specific purposes of the experiments. Ethical approval for the study was obtained from the Ethics Committee of NTT Communication Science Laboratories (approval number: R06-013). The experiments adhered to the principles of the 2013 Declaration of Helsinki. Written informed consent was obtained from all participants prior to their participation.

### Apparatus

Stimuli were presented on an LCD monitor (Display++, Cambridge Research Systems Inc., USA). The monitor’s luminance output was linearly calibrated using a luminance meter (LS-150, Konica Minolta Inc., Japan), with a luminance range of 0 to 144 cd/m^²^. A Mac Pro computer (Apple Inc., USA) was used to control stimulus presentation and data collection. The experimental scripts were written by using PsychoPy^15^ . The observation distance was 0.573 m.

### Stimuli

Stimuli were two-dimensional 1/f noise images that were sequentially deformed (Movie 1; see also Figure 1 for details of stimulus generation). Each stimulus measured 256 × 256 pixels and was presented for 0.5 seconds at a frame rate of 15 Hz. For each trial, a new 1/f noise image was generated and deformed as described below. We chose the 1/f noise as a background image based on the previous study^9^ . To deform the image, two sets of two-dimensional white noise were generated as deformation maps: one for horizontal deformation and the other for vertical deformation. Each set consisted of 15 frames, with each frame measuring 256 × 256 pixels. The white noise was filtered either using a low-pass filter (Experiment 1) or a band-pass filter (Experiments 2 and 3). For low-pass filtering, the cut-off spatial frequency was set at one of four levels: 4, 8, 16, or 32 cycles per image (cpi), corresponding to 0.4, 0.8, 1.6, and 3.2 cycles per degree (cpd). The maximum wavelengths for these levels were 2.5, 1.25, 0.63, and 0.31 degrees, respectively. For band-pass filtering in Experiment 2 (Movie 2), the range was set to one of four octave bands: [2–4, 4–8, 8–16, and 16–32 cpi], corresponding to [0.2–0.4, 0.4–0.8, 0.8–1.6, and 1.6–3.2 cpd]. The filtered noise values were standardized to a range of -12 to 12 and were used as deformation magnitudes. The deformation was implemented using OpenCV’s cv.remap function, which applied the deformation maps to the 1/f noise. For each frame, the white noise was spatially displaced in one of four directions (upward, downward, leftward, or rightward) with magnitudes of 2, 4, 8, 16, 32, 64, or 128 pixels, corresponding to 0.07, 0.15, 0.31, 0.62, 1.25, 2.5, and 5 degrees, respectively. The deformed 1/f noise was then processed with a two-dimensional Tukey window with a diameter of 9 degrees. The final stimuli were presented to participants at a frame rate of 30 Hz for 0.5 seconds. In Experiment 3, we aimed to eliminate the influence of image deformation on direction discrimination. To achieve this, the original (intact) 1/f noise was subtracted from the deformed stimuli, and the resulting subtraction images were overlaid on a uniform neutral gray background before being presented to participants (Movie 3).

### Procedure

Each participant was tested individually in a dimly lit experimental room. A head and chin rest were used to stabilize the participant’s head and ensure a consistent visual field. To initiate each trial, participants pressed the space bar. Following this, sequences of deformed 1/f noise images were presented for 0.5 seconds. After the sequence disappeared, participants reported the perceived direction of deformation-based motion using a four-alternative forced-choice (4AFC) task by pressing assigned keys. Each participant completed four sessions, with each session consisting of 140 trials. The trials included seven levels of displacement magnitude, four motion directions, and five repetitions per condition. The trial order within each session was randomized. In Experiment 1, one cut-off frequency condition was tested per session. In Experiments 2 and 3, one frequency range condition was tested per session. The order of sessions was randomized within each experiment. Each participant completed all four sessions in approximately one hour, including breaks.

### Statistics

The proportion of correct responses for discriminating the direction of deformation-based motion was calculated for each experimental condition. In Experiment 1, a two-way repeated-measures ANOVA was performed with cut-off frequency and displacement magnitude as within-participant factors. In Experiments 2 and 3, a two-way repeated-measures ANOVA was conducted with frequency bands and displacement magnitude as within-participant factors. Due to violations of the sphericity assumption, Greenhouse-Geisser corrected p-values were used for interpreting the results. For post-hoc tests, Bonferroni-corrected p-values were applied. Effect sizes were reported using eta-squared (*η^²^*).

## Results

Supplementary Data 1 provides the raw data for Experiments 1, 2, and 3.

Supplementary Data 2, 3, and 4 present the results of ANOVAs for Experiments 1, 2, and 3, respectively. Hereinafter, we summarize the significant findings of the experiments. For further details, please refer to the supplementary data.

### Experiment 1

Figure 2a shows proportion correct for the direction discrimination as a function of displacement magnitudes for each cut-off frequency condition. We conducted a two-way ANOVA and acknowledged that both the main effect of the cut-off frequency (*F*(3,42) = 13.682, *p*_*gg_corrected*_ < .0001, *η*^*2*^ = 0.14, *ε* = 0.70) and the main effect of the displacement magnitude (*F*(6,84) = 1061.001, *p*_*gg_corrected*_ < .0001, *η*^*2*^ = 0.938, *ε* = 0.69) were significant. The interaction between the two factors was significant (*F*(18, 252) = 121.913, *p*_*gg_corrected*_ < .0001, *η*^*2*^ = 0.833, *ε* = 0.22). For all conditions of cut-off frequencies, the simple main effect of the displacement magnitude was significant (*p* < .0001). The simple main effect of the cut-off frequencies was significant (*p* < .005) For all displacement magnitude, except when the displacement magnitude was 5 deg. When the cut-off frequency was 0.4 cpd, the performance peaked at the displacement magnitudes of 0.62 and 1.25 deg. That is, the performance was best at specific displacement magnitudes. On the other hand, when the cut-off frequencies were 0.8, 1.6, and 3.2 cpd, the performance was better when the displacement magnitudes were smaller. The results show that the discrimination performance as a function of the displacement magnitude depends highly on the cut-off frequencies, indicating that different mechanisms underlie the discrimination of deformation-based directions depending on the cut-off frequencies.

**Figure 2.**
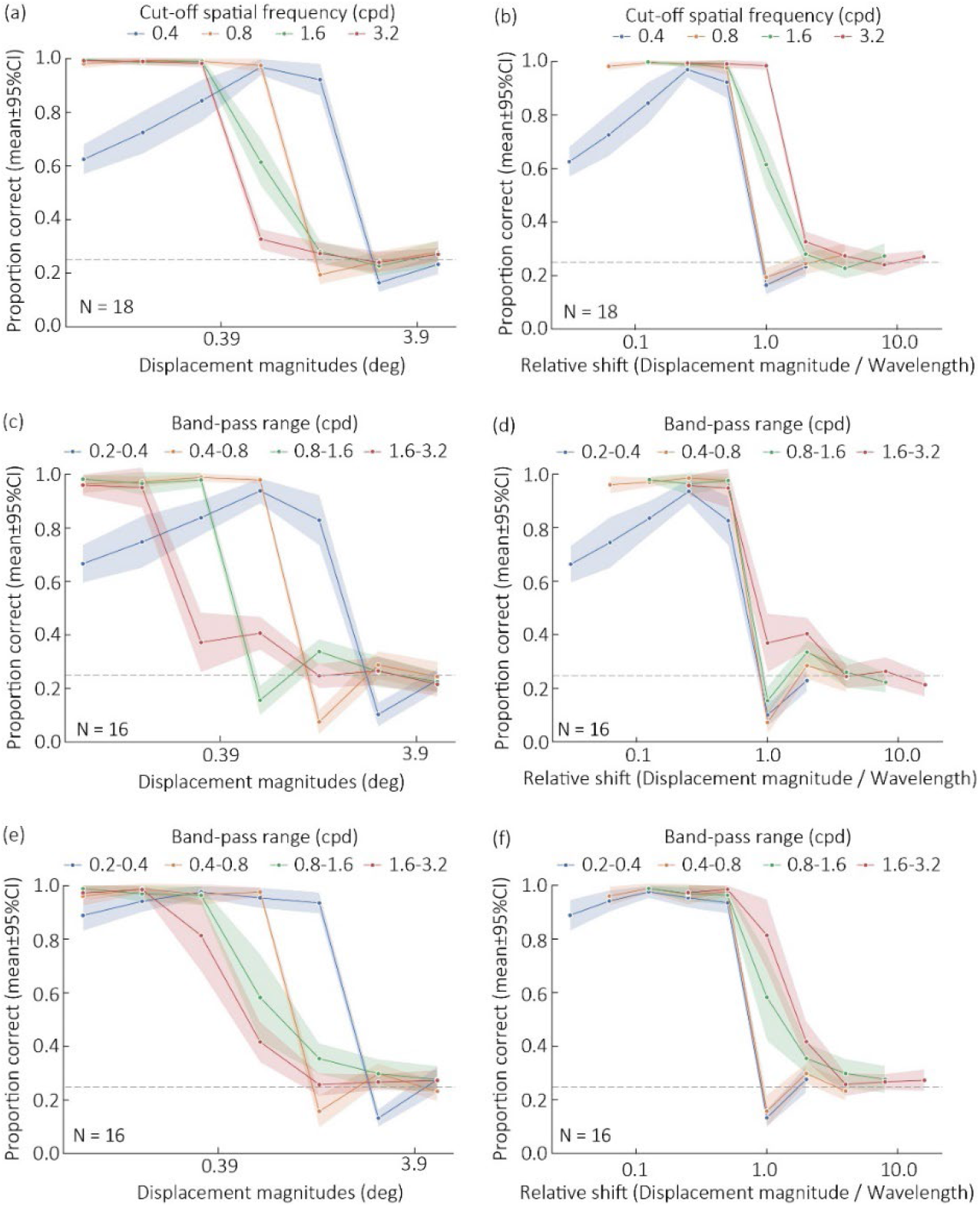
Experimental results from (a) Experiment 1, (c) Experiment 2, and (e) Experiment 3, with the results replotted as a function of relative shift in (b), (d), and (f), respectively.

Moreover, when the cut-off frequencies were lower, the discrimination performance was higher at larger displacement magnitudes. Specifically, when the displacement magnitude was 0.62 deg, the performance was significantly lowest at 3.2 cpd, but it was significantly highest at 0.4 and 0.8 cpd. These results are consistent with the previous study^14^ which showed that higher spatial frequencies resulted in a lower maximum displacement for the perception of coherent motion.

Next, we examined the proportion of displacement magnitude relative to the wavelength, referred to as the “relative shift,” as we hypothesized that analyzing this measure is essential to evaluate whether the characteristics for discerning deformation-based motion direction are consistent across different cut-off frequency conditions. Figure 2B presents the proportion of correct responses as a function of relative shift for each cut-off frequency condition. As the cut-off frequency increased, performance declined at higher relative shifts. These results suggest that the ability to discriminate motion direction is not consistent across cut-off frequencies at this stage.

One possible explanation for this inconsistency is the use of low-pass filters to generate deformation maps. In higher cut-off frequency conditions, the stimuli contain more broadband frequency components of image deformation, which may contribute to retaining motion direction discrimination even at higher relative shifts.

### Experiment 2

Experiment 2 investigated how performance declines in response to relative shifts when deformation maps are generated using band-pass filters. Instead of cut-off frequencies, we examined the role of frequency bands of band-pass filters in the direction discrimination. Specifically, we tested the following four frequency bands: 0.2-0.4, 0.4-0.8, 0.8-1.6, and 1.6-3.2 cpd. Figure 2c shows proportion correct for the direction discrimination as a function of displacement magnitudes for each frequency band condition. As in Experiment 1, we conducted a two-way ANOVA and acknowledged that both the main effect of the frequency bands (*F*(3,45) = 27.686, *p*_*gg_corrected*_ < .0001, *η*^*2*^ = 0.26, *ε* = 0.77) and the main effect of the displacement magnitude (*F*(6,90) = 641.944, *p*_*gg_corrected*_ < .0001, *η*^*2*^ = 0.885, *ε* = 0.53) were significant. The interaction between the two factors was significant (*F*(18, 270) = 110.446, *p*_*gg_corrected*_ < .0001, *η*^*2*^ = 0.79, *ε* = 0.26). For all conditions of frequency bands, the simple main effect of the displacement magnitude was significant (*p* < .0001). The simple main effect of the frequency bands was significant (*p* < .0001) for all displacement magnitude, except when the displacement magnitude was 5 deg (p = .855). When the frequency band was 0.2–0.4 cpd, performance peaked at a displacement magnitude of 0.62 degrees. Consistent with Experiment 1, the best performance occurred at specific displacement magnitudes. In contrast, for frequency bands of 0.4–0.8, 0.8–1.6, and 1.6–3.2 cpd, performance improved as displacement magnitudes decreased. Figure 2D illustrates the proportion of correct responses as a function of relative shift. Compared to Experiment 1, the variability in performance across frequency bands was reduced. These results suggest that when the frequency band of image deformation is properly controlled, the discrimination of deformation-based motion is processed more consistently across frequency bands.

An unresolved issue is why performance declines when the displacement magnitude is small in the lower frequency band condition (0.2–0.4 cpd). We hypothesize that, in this frequency band, the motion signal associated with image deformation itself may be stronger than or comparable to the motion signals associated with deformation-based motion. Figure 3 illustrates the pixel slices of the clip used in Experiment 2. In the lowest frequency band condition (0.2–0.4 cpd), the motion signal derived from deformation-based motion is extremely weak (Figure 3a). Simultaneously, a subtle motion signal exists in the direction orthogonal to the deformation-based motion, originating from local deformation itself (Figure 3b). This interference may prevent the visual system from accurately detecting the motion direction in the displacement direction of deformation-based motion. Conversely, in the highest frequency band condition, the motion signal from deformation-based motion is strong (Figure 3c), while the motion signal originating from local deformation is minimal or indistinct (Figure 3d). This likely enhances the detection of the motion signal from deformation-based motion.

**Figure 3.**
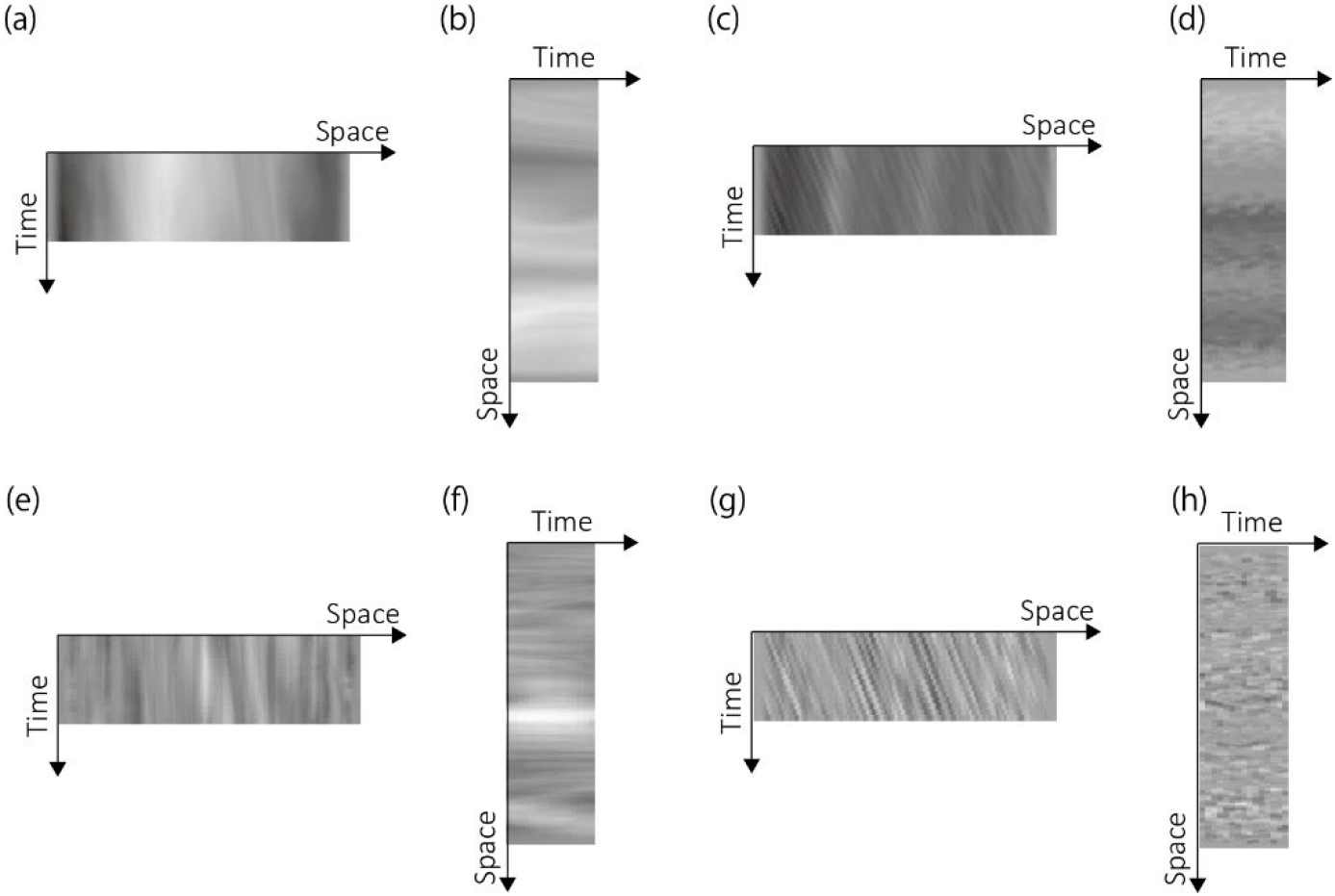
Horizontal and vertical slices of pixels in video clips used as (a-d) Experiment 2 and (e-h) Experiment 3 stimuli. (a) Horizontal and (b) vertical slices of pixels in the 0.2-0.4 cpd condition with the displacement magnitude of 0.07 deg. (c) Horizontal and (d) vertical slices of pixels in the 1.6-3.2 cpd condition with the displacement magnitude of 0.07 deg. (e) Horizontal and (f) vertical slices of pixels in the 0.2-0.4 cpd condition with the displacement magnitude of 0.07 deg. (c) Horizontal and (d) vertical slices of pixels in the 1.6-3.2 cpd condition with the displacement magnitude of 0.07 deg.

To test this hypothesis, we conducted an experiment to determine whether reducing deformation-related components in the stimuli could improve the detection of deformation-based motion signals even in the lowest frequency band condition.

#### Experiment 3

In this experiment, we subtracted an intact (non-deformed) background image from each video frame in stimulus clips as used in Experiment 2. By doing so, we could effectively attenuate the contribution of image deformation to the direction discrimination of deformation-based motion. We asked the same task to the participants and checked whether the performance did not decline in the smaller displacement magnitude in the low frequency band of image deformation.

Figure 2e shows proportion correct for the direction discrimination as a function of displacement magnitudes for each frequency band condition. As in Experiment 2, we conducted a two-way ANOVA and acknowledged that both the main effect of the frequency bands (*F*(3,45) = 40.293, *p*_*gg_corrected*_ < .0001, *η*^*2*^ = 0.232, *ε* = 0.56) and the main effect of the displacement magnitude (*F*(6,90) = 678.307, *p*_*gg_corrected*_ < .0001, *η*^*2*^ = 0.894, *ε* = 0.542) were significant. The interaction between the two factors was significant (*F*(18, 270) = 46.212, *p*_*gg_corrected*_ < .0001, *η*^*2*^ = 0.646, *ε* = 0.135). For all conditions of frequency bands, the simple main effect of the displacement magnitude was significant (*p* < .0001). The simple main effect of the frequency bands was significant (*p* < .0001) for all displacement magnitude, except when the displacement magnitude was 5 deg (*p* = .855). Importantly, in the lowest frequency band condition, the performance at 0.07 deg was statistically comparable to that at 0.15, 0.31, 0.62, and 1.25 deg (*p* > .15). Similarly, the performance at 0.15 deg was comparable to that at 0.31, 0.62, and 1.25 deg (*p* > .12), and the performance at 0.31 deg was comparable to that at 0.62 and 1.25 deg (*p* = 1.00). These results support our hypothesis that discrimination of deformation-based motion direction does not decline with smaller displacement magnitudes in the lowest frequency band condition.

Figure 2f shows the proportion of correct responses as a function of the relative shift. Compared to the results from Experiment 2 (Figure 2d), the discrimination performance was less consistent across different frequency band conditions, particularly at larger displacement magnitudes. We speculate that this inconsistency may be due to the subtraction of the original (intact) background image from the deformed images, which likely reduced interference from local deformations in the discrimination of deformation-based motion. However, the exact source of this interference remains an open question.

## Discussion

The present study investigated the properties of discriminating the direction of deformation-based motion. Experiments 1 and 2 demonstrated that the spatial frequency components of image deformation are key determinants of direction discrimination in terms of displacement magnitude. Specifically, lower spatial frequencies of image deformation contributed to maintaining discrimination performance even at larger displacement magnitudes. Experiment 3 further revealed that attenuating local deformation enhanced discrimination performance, particularly at lower spatial frequencies, suggesting a potential interaction between the processing of deformation-based motion and local deformation signals.

Our findings indicate that the discrimination of deformation-based motion direction relies on linear motion detection, which is driven by the displacement of deformation patterns. As shown in Figures 3c and 3g, the displacement of image deformation generates spatiotemporally tilted orientations across frames. These spatiotemporal orientations serve as a robust cue for the human visual system to detect motion^16^ . Even in the context of deformation-based motion, such fundamental motion detection mechanisms operate effectively.

However, our results also suggest a connection between the detection of motion signals arising from the displacement of image deformation and those originating from local deformation. In Experiment 3, eliminating local deformation enhanced discrimination performance, particularly in the lower spatial frequency bands. Interestingly, participants could perceive both the direction of deformation-based motion and the local deformation simultaneously in the stimuli used in Experiments 2 and 3. Therefore, future studies should investigate how the discrimination of deformation-based motion direction integrates with the phenomenological experience of the stimuli.

Our findings suggest that spatiotemporal orientation signals derived from deformation-based motion play a crucial role in the perception of transparency in image deformation. As shown in Figure 3c, two distinct types of spatiotemporal orientations are evident: one arising from the linear shift of luminance across frames, and the other from vertical luminance contrasts associated with static textures. This indicates that perceptual transparency in image deformation is, at least partially, facilitated by the segregation of dynamic luminance flow from static luminance signals.

In this study, we used 1/f noise as the background, but we would like to discuss its influence. We selected 1/f noise as the background image because its contrast characteristics are similar to those of natural images^17,18^ . However, elements like riverbed pebbles or grass create sharp contrast edges in retinal images. Therefore, conducting follow-up experiments with backgrounds that have clear luminance edges would be important for elucidating the mechanisms of deformation-based motion perception. That said, it is worth noting that the previous study^9^ has demonstrated that dynamic image deformation contributes to transparency perception, regardless of whether the background consists of natural images with sharp edges or 1/f noise images.

Local deformation has been discussed in this study as a factor that hinders the perception of deformation-based motion direction. However, in real-world contexts, the perception of local deformation likely plays an important role in material perception. For example, the degree of local deformation could serve as a cue for estimating the thickness of a transparent medium^19^ . Thus, the direction of deformation-based motion and the degree of local deformation are likely perceived simultaneously. While this study focused exclusively on directional perception, future research could investigate how humans integrate material perception based on local deformation with the directional perception of deformation-based motion in natural environments. Such investigations would provide valuable insights for the perception of materials from image deformation.

## Acknowledgements and funding statement

We would like to thank Miyuki Otani for her assistance with data collection during the experiments. This study received no specific funding.

## Notes

### Competing Interest Statement

The authors have declared no competing interest.

